# A microfluidic strategy to capture antigen-specific high affinity B cells

**DOI:** 10.1101/2023.07.12.548739

**Authors:** Ahmed M. Alhassan, Venktesh S. Shirure, Jean Luo, Bryan B. Nguyen, Zachary A. Rollins, Bhupinder S. Shergill, Xiangdong Zhu, Nicole Baumgarth, Steven C. George

## Abstract

Assessing B cell affinity to pathogen-specific antigens prior to or following exposure could facilitate the assessment of immune status. Current standard tools to assess antigen-specific B cell responses focus on equilibrium binding of the secreted antibody in serum. These methods are costly, time-consuming, and assess antibody affinity under zero-force. Recent findings indicate that force may influence BCR-antigen binding interactions and thus immune status. Here, we designed a simple laminar flow microfluidic chamber in which the antigen (hemagglutinin of influenza A) is bound to the chamber surface to assess antigen-specific BCR binding affinity of five hemagglutinin-specific hybridomas under 65- to 650-pN force range. Our results demonstrate that both increasing shear force and bound lifetime can be used to enrich antigen-specific high affinity B cells. The affinity of the membrane-bound BCR in the flow chamber correlates well with the affinity of the matched antibodies measured in solution. These findings demonstrate that a microfluidic strategy can rapidly assess BCR-antigen binding properties and identify antigen-specific high affinity B cells. This strategy has the potential to both assess functional immune status from peripheral B cells and be a cost-effective way of identifying individual B cells as antibody sources for a range of clinical applications.

## INTRODUCTION

Activation, clonal expansion, and affinity maturation of B cells in germinal centers are considered the hallmarks of adaptive immunity, which are triggered when challenged by foreign antigens (e.g., viral or bacterial infection) ^1^. Upon re-challenge, memory B cells (B_mem_) differentiate rapidly to plasma cells which then secrete antigen-specific high-affinity antibodies to facilitate rapid pathogen clearance ^2^. The serum antibody pool largely reflects the antibody-producing plasma cell population, and thus does not necessarily reflect the dynamic characteristics and immunogenic potential of antigen-specific B_mem_, which are known to arise earlier from germinal center responses than plasma cells and thus carry overall fewer mutations^3^. The ability to precisely measure B cell-antigen interaction strength through the membrane bound B cell receptor (BCR) would thus greatly enhance the evaluation of functional immunity (i.e., the immune status of an individual following infection or vaccination) as it may better correlate with immune protection ^4^. Moreover, the ability to isolate antigen-specific B cells with known antigen-binding avidities could aid in rapid identification and creation of monoclonal antibody-based therapeutics.

Direct measurements of antibody-secreting cells can be performed using ELISPOT, or more recently, by flow cytometry (FACS) with surface markers and fluorescently labeled antigens ^5–7^. The latter technique can also be used to isolate antigen-binding B cells, but it does not control the force of interaction between the cell and antigen. More specifically, the fluorescent conjugated antigens are simply incubated with the cells, allowed to come to equilibrium, and the amount of the probe bound to cells, which is proportional to the affinity, is used to separate the cells. *In vivo*, B cells interrogate antigens with the B cell receptor (BCR) under force, and naïve B cells are strongly activated when this force exceeds 50 pN ^8^. This force may modify a range of B cell responses, including activation and antigen internalization ^9^. Furthermore, overall B cell binding avidity (sometimes referred to as “effective affinity”) depends on both epitope density and the intrinsic affinity of the BCR to the cognate antigen.

While BCR binding affinity is generally acknowledged to be the primary determinant of B cell activation and recruitment *in vivo* and thus as prognostic of immune protection ^10, 11^, B cells recruited to the germinal center generally encounter the same epitope density, and thus intrinsic affinity of the BCR is a useful surrogate. There are currently no tools to accurately test the overall breadth of membrane-bound BCR affinity or avidity at the single cell or the population level. Developing these tools could provide more reliable methods to monitor an individual’s immune protection status and thus could enhance vaccination strategies (e.g., distribution, volume, frequency) against existing and future infectious agents.

Finally, it is important to recognize that antibodies having similar affinity (ratio of kinetic on-rate and off-rate) can have on- and off-rates that vary over four orders of magnitude ^12^. The kinetic on- and off-rates themselves can impact B cell biology. Indeed, antibody maturation and selection, at least in some cases, has been ascribed to enhancement of the kinetic on-rate ^13, 14^. While there are established techniques available to characterize secreted antibodies, such as surface plasmon resonance, no tools are available for the measurement of kinetic properties of the cell membrane-bound BCR.

Utilizing a simple microfluidic strategy to control shear force and antigen presentation, we have developed a method to capture and enrich antigen-specific high affinity B cells, and quantify force-dependent B cell binding avidity and kinetic properties for a pool of B cells. Our results demonstrate that 1) both (shear) force and bound lifetime can be used to enrich a population of antigen-specific high affinity B cells, and 2) the affinity constants of the B cells measured in the device correlate well with the affinity of the secreted antibodies measured in solution.

## MATERIALS AND METHODS

### Mice

C57BL/6 mice (The Jackson Labs) and “SwHEL” BCR transgenic mice expressing a BCR specific for hen egg lysosome (HEL) ^15^ were provided with food and water at libitum and held under SPF housing conditions at the University of California (UC) Davis. Breeding pairs for SwHEL mice were obtained from Dr. Roger Sciammas (UC Davis) with kind permission from Dr. Robert Brink (Garvan Institute of Medical Research, New South Wales, Australia). Male and female mice, age 8 – 15 weeks were used as the source of HEL-specific B cells. All experiments involving mice were conducted in strict adherence to protocols approved by the UC Davis Animal Care and Use Committee.

### Hybridomas

To assess adherence of influenza hemagglutinin-specific B cells, we assessed five previously characterized hybridoma cell lines generated from influenza A/8/34 immunized BALB/c mice. As a negative control hybridoma we used DS.1, specific to IgMa (**Table 1**). Cells were grown in RPMI 1640 containing 10% FBS and 1:100 P/S (Gibco™ 15140122). Cells were collected when they were about 70% confluent and looked round, smooth and quite large.

**Table 1.**
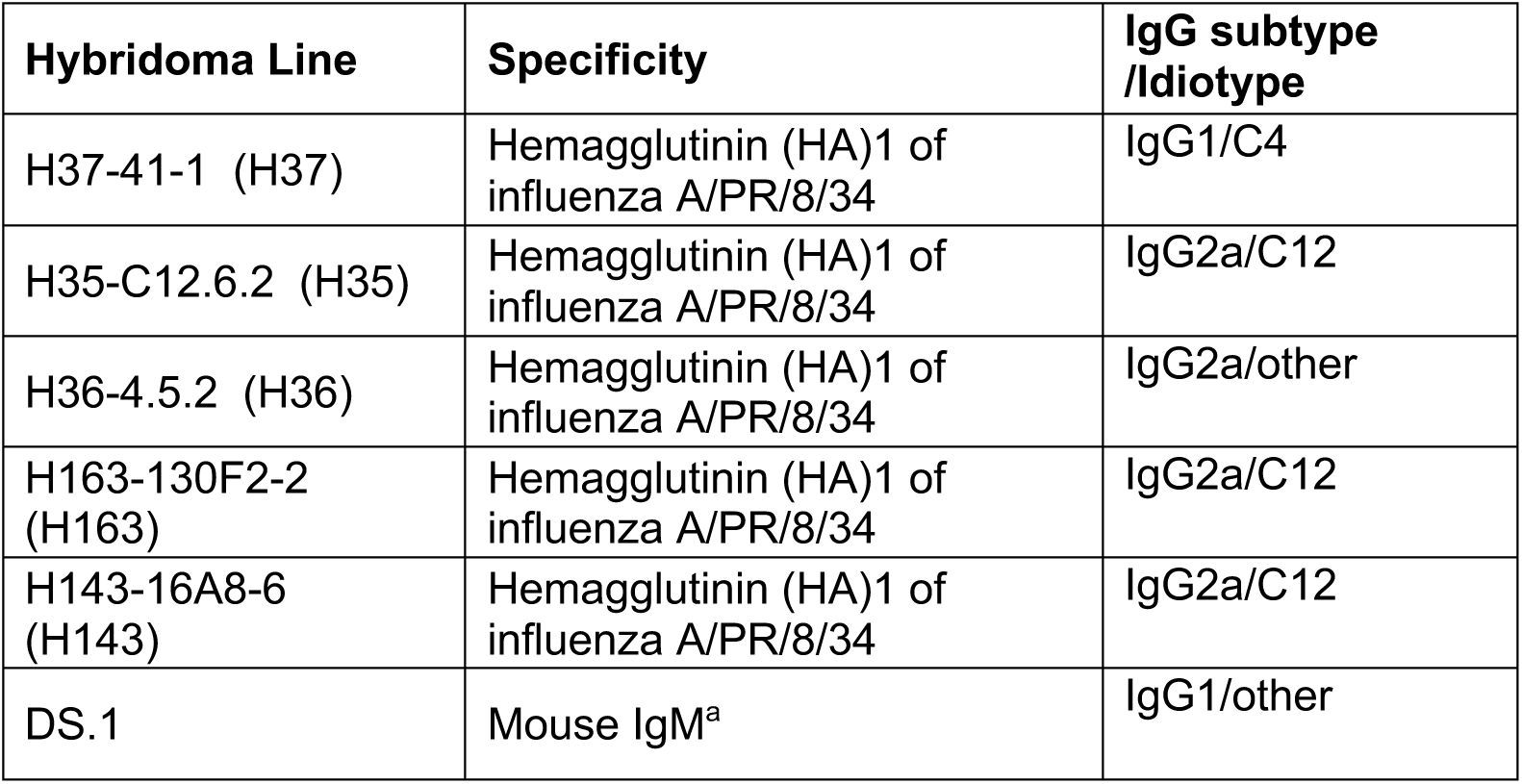
Specificity and subtype of hybridoma lines used in experiments.

| Hybridoma Line | Specificity | IgG subtype /Idiotypic |
| --- | --- | --- |
| H37-41-1 (H37) | Hemagglutinin (HA)1 of influenza A/PR/8/34 | IgG1/C4 |
| H35-C12.6.2 (H35) | Hemagglutinin (HA)1 of influenza A/PR/8/34 | IgG2a/C12 |
| H36-4.5.2 (H36) | Hemagglutinin (HA)1 of influenza A/PR/8/34 | IgG2a/other |
| H163-130F2-2 (H163) | Hemagglutinin (HA)1 of influenza A/PR/8/34 | IgG2a/C12 |
| H143-16A8-6 (H143) | Hemagglutinin (HA)1 of influenza A/PR/8/34 | IgG2a/C12 |
| DS.1 | Mouse IgM<sup>a</sup> | IgG1/other |

### B cell isolation

For binding studies involving primary B cells, cells were obtained from spleens of SwHEL or wildtype C57BL/6 mice. The study was approved by the IACUC at the University of California, Davis. Single cell suspensions were generated by grinding spleens between the frosted ends of two glass slides and filtered through a 70mm nylon mesh. All samples were then treated with ACK lysis buffer ^16^, re-filtered through nylon mesh, and resuspended in RPMI or staining media. Single cell suspensions were counted and blocked with anti-FcγR (mAb 2.4.G2). Then cells were labeled with biotinylated HEL generated in-house and anti-mouse CD19-CF594 (ID3, Biolegend), followed by staining with streptavidin-allophycocyanin and live/dead Aqua (Thermo Fisher) to assess frequencies of HEL-binding cells by flow cytometry, which were about 20-25% of total cells.

### Flow cytometry

For staining of surface Ig, hybridoma cells were collected, washed and resuspended in staining buffer. Cells were blocked with anti-FcgR (2.4.G2), surface stained with APC-Cy7 anti-mouse Ig kappa light chain (BD 561353) then stained with Live/Dead Aqua (Thermo Fisher, L34966). Each step was done for 15 min on ice followed by washing the cells in staining buffer. Cells were analyzed using a BD FACS Symphony flow cytometer. Data analysis was done using FLowJo software.

### Purification of monoclonal antibodies

Hybridoma supernatant was collected 3-5 days after cells were seeded into flasks when medium turned from pink to orange and filtered through a 0.22 µm filter followed by ammonium sulfate precipitation using ammonium sulfate salt, added slowly (313.5g to 1000ml) to reach ∼50% saturation and then incubated for 5–15 h at 4°C. Supernatants were centrifuged at 5000*g* for 30 min at 4°C. The pellet was resuspended in PBS and then dialyzed against at least three changes of PBS for 24–48 h. IgG was purified by low pressure, HiTrap Protein G column chromatography following the manufacturer’s instructions (Cytica HiTrap Protein G HP, 17040501). After elution, antibody was concentrated and buffer exchanged into PBS or storage buffer (10 mM Tris, 150 mM NaCl, 0.1% NaN3, pH 8.2). The protein concentration was determined by measuring the optical density at 280 nm. For IgG, an absorption of *OD*_280_ of 1.35 was set to equal 1 mg/mL IgG.

### ELISA

Influenza virus-specific ELISA was performed as described ^16^. Briefly, ELISA plates (MaxiSorp 96 well plates, Thermo Fisher #12-565-135) were coated overnight at room temperature with influenza A/Puerto Rico/8/34 virus particles (400 HAU/ml; in house) purified from the allantoic fluid of infected day 14 embryonated hen eggs, precipitated with polyethylene glycol and purified via a sucrose gradient centrifugation. Plates were washed and non-specific binding was blocked with 1% newborn calf serum, 0.1% dried milk powder, and 0.05% Tween 20 in PBS (ELISA blocking buffer). Following pilot studies, all HA-specific mAb were added to the plate at a starting dilution of 100 ng/ml (except H143-16A8-6 which was used at 10 μg/ml) then were serially diluted by two-fold increments in PBS. Binding was revealed with biotinylated anti-IgG (Southern Biotech 1030-08), followed by streptavidin horseradish peroxidase (Vector SA-5004) both diluted in ELISA blocking buffer. Substrate (0.005% 3,39,5,59-tetramethylbenzidine in 0.05 M citric acid buffer, PH 4.0 and 0.015% hydrogen peroxide (Spectrum H1070). The reaction was stopped with 1N sulfuric acid after 20 minutes. Absorbance was measured at 450 nm (595 nm reference wavelength) on a spectrophotometer (SpectraMax M5, Molecular Devices).

### Microfabrication and device surface preparation

Microfluidic devices were prepared using standard methods of soft lithography ^17, 18^. In brief, a SU8 master mold was prepared and the microdevice was created by casting polydimethylsiloxane (PDMS; Dow Corning, Midland, MI) on the SU-8 master mold. Once, polymerized, the PDMS was peeled off the master mold. Glass slides (Thermo Fisher Scientific) were rinsed in purified water and were then plasma bonded to the PDMS to form channels of 10 mm x 0.8 mm x 0.1 mm (length x width x height).

The microfluidic devices were coated with the desired antigens following well documented streptavidin-biotin chemistry ^19^. In brief, the devices were incubated at room temperature with (3-Mercaptopropyl)trimethoxysilane (3.65% in absolute ethanol, Sigma-Aldrich) for 1 hour, washed twice with absolute ethanol, and then incubated with N-γ-maleimidobutyryl-oxysuccinimide ester (1mM in absolute ethanol, Thermo Fisher Scientific) for 30 min, washed twice with absolute ethanol and finally incubated with NeutrAvidin at 4°C (100 ug/mL in PBS, Thermo Fisher Scientific) for 2 days, washed twice with PBS. The devices were then incubated at 4°C with the biotinylated antigen prepared in PBS at a desired concentration for 2 days, washed twice with PBS and incubated with BSA (10 mg/mL in PBS) for 15 minutes before perfusing the cells.

### Cell perfusion through the microfluidic device

We used a pipette tip (200 μl) connected at the inlet as the entry port for cells and fluid, and the outlet of the microfluidic chip was connected by Tygon tubing to a high precision syringe pump (Harvard Apparatus), operated in withdraw mode. The devices were first equilibrated by perfusing PBS for 2 min at 200 μl/min . The devices were placed on an IX83 inverted microscope (Olympus) equipped with a high-speed camera and hardware to acquire stream acquisitions.

Cells suspended at a prescribed concentration (generally 300,000 cells/mL) were added to the source tip after setting desired flow. The devices were imaged using a brightfield, 10X objective, and an image acquisition speed of 8 frames per second. Four different flows were used (10, 20, 50, and 100 μL/h) corresponding to wall shear stress of 0.03, 0,06, 0,15, and 0.3 dynes/cm^2^ or tensile stress of the bond of approximately 65, 130, 325, and 650 pN ^20^. For higher flows (50 and 100 μL/h), the acquisition speed was increased to 30 frames per second, and, to accommodate for this higher acquisition speed, total acquisition time was reduced to 2 minutes. The images were saved for later analysis.

In some experiments, the hybridoma cell lines were labelled with either CellTracker BMQC (violet), CMFDA (green), CMMTR (Orange), or Deep Red. A mixture of differently labelled hybridoma cells at 1.5 x 10^6^ cells/mL was perfused through the device. The devices were imaged using a FV1200 Fluoview confocal laser scanning microscope (Olympus) connected to FV10-ASW image acquisition and analysis software (Olympus). The confocal microscope was used for simultaneous time lapse recording of four color channels. The images were acquired at the maximum allowable acquisition rate of ∼1 frame per second.

### Analysis of cell binding in the device

Each cell perfusion experiment was analyzed to find: 1) total number of cells flowing across the field of view; 2) the number of cells bound to the surface; and 3) the time each cell remained bound (bound lifetime). To find these, the microscopy-obtained time lapse image sequences were analyzed using TrackMate (an ImageJ plugin) following the recommended protocol ^21^. The software records cell positions from each image in a time sequence. The algorithm then generates tracks of cell movements using consecutive images from the sequence. The software generates images with tracks that are color-coded by the order in which the cells enter the field of view. The “tracks” and “spots” data files generated from TrackMate were exported to the RLNEK (Receptor-Ligand Non-Equilibrium Kinetics) ^22^ to compute capture efficiency and bound lifetimes. Capture efficiency is defined as the ratio of the number of bound cells (N_b_) to the total number of cells (N_0_) flowing across the field of view. A binding event was defined as a cell moving less than 0.5 µm. The minimum binding time criterion (i.e., cells remain bound for greater than this time) was chosen as 10, 20, 50 or 100 s.. In a small number of cases where a mixture of cells was perfused, the acquisition speed was < 5 frames per second and an accurate count with the automated analysis was not possible; in these cases, the analysis was performed manually.

### Mathematical model of cell perfusion through the microfluidic device

A mathematical model capable of simulating laminar fluid flow and cell movement through the device was created using COMSOL Multiphysics® 5.2a software. The computer aided design (CAD) file of the device with all 3-dimenstions was imported as the geometry of the device. The laminar flow module was used to drive flow through the device at a desired fluid flow rate. The no-slip boundary condition was applied for all surfaces except the microfluidic entrance and exit. To find the cell trajectories, Particle Tracing Module was used. Particle properties were set as typical cell properties (radius = 5 μm; r = 1.086 g/cc). The cells entering the device through the inlet and their initial position at the inlet boundary were set at randomly chosen locations. The model was set so that cells entered the device in a short pulse of 1s and at every 10s interval thereafter. The number of cells entering every 10s with each pulse were computed using the known perfusion concentration of the cells (300,000 cells/mL) and Q = 10, 20, 50, 100, 200 μL/h. Because of gravity, the cells eventually settled in the device. They also experienced fluid drag under the laminar fluid flow, which moved them along the length of the device. The cells that first contact the bottom surface of the device and the cells exiting the device were frozen at those boundaries to create a visual demonstration of the simulations.

### Estimation of Effective 𝒌_𝒐𝒏_ and 𝒌_𝒐𝒇𝒇_

Assuming cell binding to the substrate as a first-order process, the number of bound cells should obey the following rate law : 𝑑𝑁_*b*(1)_⁄𝑑𝜏 = 𝑘_*on*_(𝑁_*_ − 𝑁_*b*(1)_), where 𝜏 is time that the cell needs to pass through the field of view or residence time of cells flowing across the field of view, 𝑁_*b*(1)_ is the number of cells that bind to form the initial tether (or bind at least 1 s), and 𝑁_*_ is the total number of cells that enter the field of view per 𝜏. The analytical solution to the above equation is: 𝑘_*on*_ = (−ln ((𝑁_*_ − 𝑁_*b*(1)_)⁄𝑁_*_)⁄𝜏 . We performed simple linear regression on the data to determine 𝑘_*on*_ ^23^. Thus, 𝑘_*on*_ determined in our assay is a function of number and diffusivity of BCR/ligands on the substrate and the intrinsic single molecule on-rate.

A bound cell detaches from the substrate due to kinetic off-rates. We assumed a first order process for cell detachment: 𝑑𝑁_*b*_⁄𝑑𝑡_*b*_ = 𝑘_*off*_𝑁_*b*_, where 𝑁_*b*_ is the number of bound cells for at least time 𝑡_*b*_. The analytical solution for this equation is: 𝑘_*off*_ = ln (𝑁_*b*(*tb*)_⁄𝑁_*b*(10)_) /(𝑡_*b*_ − 10) where 𝑁_*b*(*tb*)_ and 𝑁_*b*(10)_ are the number of cells that bind at least t_b_ and 10 s, respectively, Thus, the off-rate is simply the slope of the line graph between fraction of bound cells and their bound lifetime, which is also referred to as survival curves ^24, 25^. Data fits were weighted by the fraction of bound cells. The force-dependent 𝑘_*off*_ is related to force independent *k_off_^0^* by the Bell model ^26^, 𝑘_*off*_ = *k*^0^_*off e^XβF/KBT^*_, where 𝑥_β_ is reactive compliance of the intermolecular bond, F is the force applied on the bond, and 𝐾_*B*_ is Boltzmann’s constant. In the present study, F is varied by applying fluid shear force (F can be calculated from the wall shear stress as detailed previously ^22^) and linear regression was performed on log transformed data to determine *k*^0^_*off*_.

### Oblique-Incidence Reflectivity Difference

Five purified anti-HA IgG molecules, H35-c12.6.2, H36-4.5.2, H37-41-1, H143-12, and H163-12-2 (Table 1) were separately diluted with 1× PBS to printing concentration of 6.7 µM (i.e. 1 mg/ml). Bovine serum albumin (BSA) and biotinylated bovine serum albumin (BBSA) were diluted separately with 1× PBS to printing concentration of 3.8 µM (i.e. 0.25 mg/ml). On an epoxide-functionalized glass slide (1”×3”), we printed 6 identical microarrays from these 8 printing solutions. Each array consists of 39 replicates of 5 IgG molecules and BSA, and 3 replicates of BBSA. BSA and BBSA are negative and positive controls, respectively. Each microarray was housed in a separate, optically accessible reaction chamber (12 mm × 6 mm × 0.4 mm; i.e., volume = 29 µL). Before binding assays were conducted, the microarrays were washed with 1× PBS, blocked with a solution of BSA at 2 mg/mL in 1× PBS for 30 minutes, and then washed again in 1× PBS.

For affinity binding assays, we prepared 300 nM solutions of influenza A/PR/8/34 virus H1N1 haemagglutinin (HA) recombinant antigen (or Rec-HA; The Native Antigen Company, Kidlington, Oxfordshire, UK) in 1x PBS. For avidity binding assays, we prepared a 2×10^-4^ HAU/ml solution of influenza A/PR/8/34.

For the affinity binding reactions, we first replaced the 1x PBS in the reaction chamber with the HA solution and then incubated the microarray in the HA solution under a slow flow condition at 2.5 µL/min for 30 minutes for the association phase of the reaction. After the association phase, we replaced the HA solution in the chamber with 1x PBS and then incubated the microarray in 1× PBS (under a slow flow condition at 20 µL/min) for another 90 minutes for the dissociation phase of the reaction.

For avidity binding reaction, we replaced the 1x PBS in the reaction chamber with a solution of 2 x 10^-4^ HAU/ml A/PR8 and then incubated the microarray in this solution under a slow flow condition at 2.1 µL/min for 4 hours for the association phase. After the association phase, we replaced the A/PR8 solution with 1x PBS and then incubated the microarray again under a slow flow condition at 10 µL/min for another 2 hours for the dissociation phase of the reaction.

To measure binding curves during the reaction, we used an oblique-incidence reflectivity difference (OI-RD) scanner, described previously ^12, 27–29^. With this scanner, we measured the phase change in a reflected optical beam due to the presence of a biomolecular layer on a solid support during the reaction. Similar to a surface-plasmon-resonance (SPR) sensor, the phase change detected with an OI-RD scanner is converted to the surface mass density of the biomolecular layer. In the present work, the scanner measured in real time the amount of Rec-HA or A/PR8 virions captured by printed (i.e., immobilized) IgG molecules and the control molecules. The association-dissociation curves (i.e., binding curves) were fit to yield rate constants k_on_ and dissociation rate constants k_off_. Before and after each reaction, we also acquired OI-RD images for endpoint analysis.

### Spleen Cell Labelling

SwHEL spleen cells were perfused in a device coated with HEL and then washed with PBS. Cells bound in the device were labelled with anti-mouse CD19 Alexa Fluor 647 and anti-mouse CD3 Alexa Fluor 488 antibodies. Cells in the device were imaged using a FV1200 Fluoview confocal laser scanning microscope (Olympus) connected to FV10-ASW image acquisition and analysis software (Olympus).

### Statistics

Unless otherwise mentioned, statistical significance indicates p < 0.05 by one-way ANOVA with Tukey’s multiple comparison test.

## RESULTS

To examine BCR-antigen interactions under force, we utilized a simple single rectangular-shaped microfluidic device. The glass surface was functionalized by incubating biotinylated antigen withthe surface precoated with NeutrAvidin (Fig. 1A, S1A). When cells are perfused in the device, antigen-specific cells can bind to the surface (Fig. 1B). A COMSOL Multiphysics model simulates cells entering the device and flowing in a horizontal trajectory as shown with the streamlines (Fig. 1C). To interact with the antigen, cells must be close enough to the surface to do so. The model shows the distance along the channel where the cells settle to the bottom and contact the surface at a given flow (Fig. 1D). At 100 µL/h, the highest flow at which we perfused cells, all cells settle to the surface before reaching the outlet.

**Figure 1.**
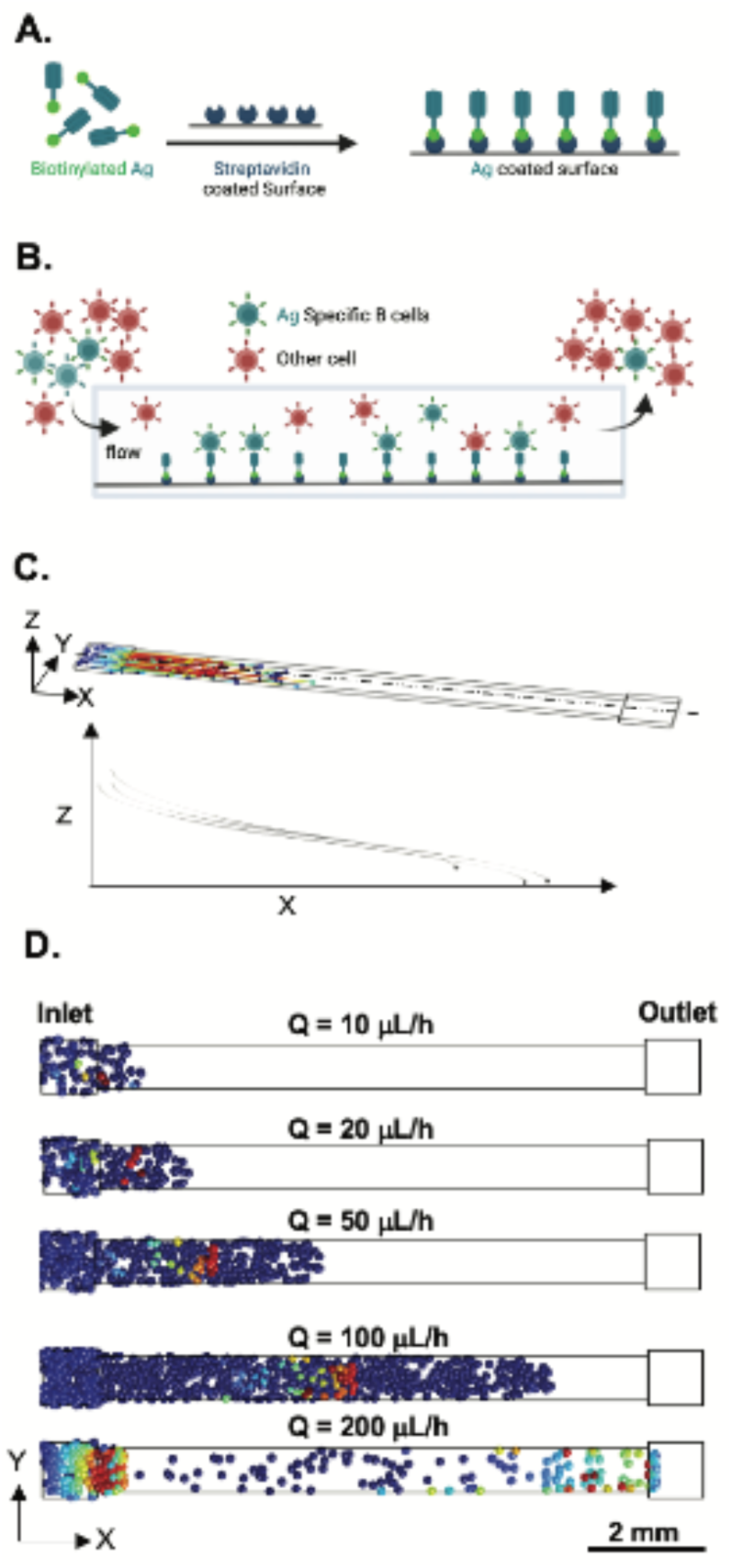
A) Schematic of antigen functionalized on the surface of the device and B) cells binding to the antigen-coated surface. C) COMSOL Multiphysics model of cells entering the microfluidic device and flowing horizontally and D) settling to the bottom along the length of the device at different flow rates. Cells that contact the surface (blue) are frozen in place.

To determine the optimal flow and antigen coating conditions for capturing antigen-specific B cells, we used high affinity HEL-specific B cells, which were harvested from genetically modified SwHEL mouse spleens (Fig. 2A). Roughly 20% of cells in the SwHEL spleen are HEL-specific B cells, as assessed by flow cytometry conducted prior to each experiment (Fig. 2B). We tested various concentrations of HEL coating on the surface. At 0.02 μM there was no measurable difference between binding of SwHEL cells in comparison to cells from wild-type (WT) mice (Fig. 2C). An increase in HEL coating concentration by an order of magnitude resulted in a clear difference in binding between WT and SwHEL cells (Fig. 2C), but additional increases in HEL concentration did not further increase cell binding. As such, an antigen coating concentration of 0.2 μM was used for all subsequent studies. Capture efficiency (∼ 20%) was comparable to flow cytometry data showing the fraction of positive cells in the WT and SwHEL spleen cell mixtures (Fig. 2C). Microscopic imaging illustrates the difference in capture between WT and SwHEL cells at different coating concentrations (Fig. 2D). All bound cells were CD19^+^/CD3^-^ consistent with B cells (Fig. S1).

**Figure 2.**
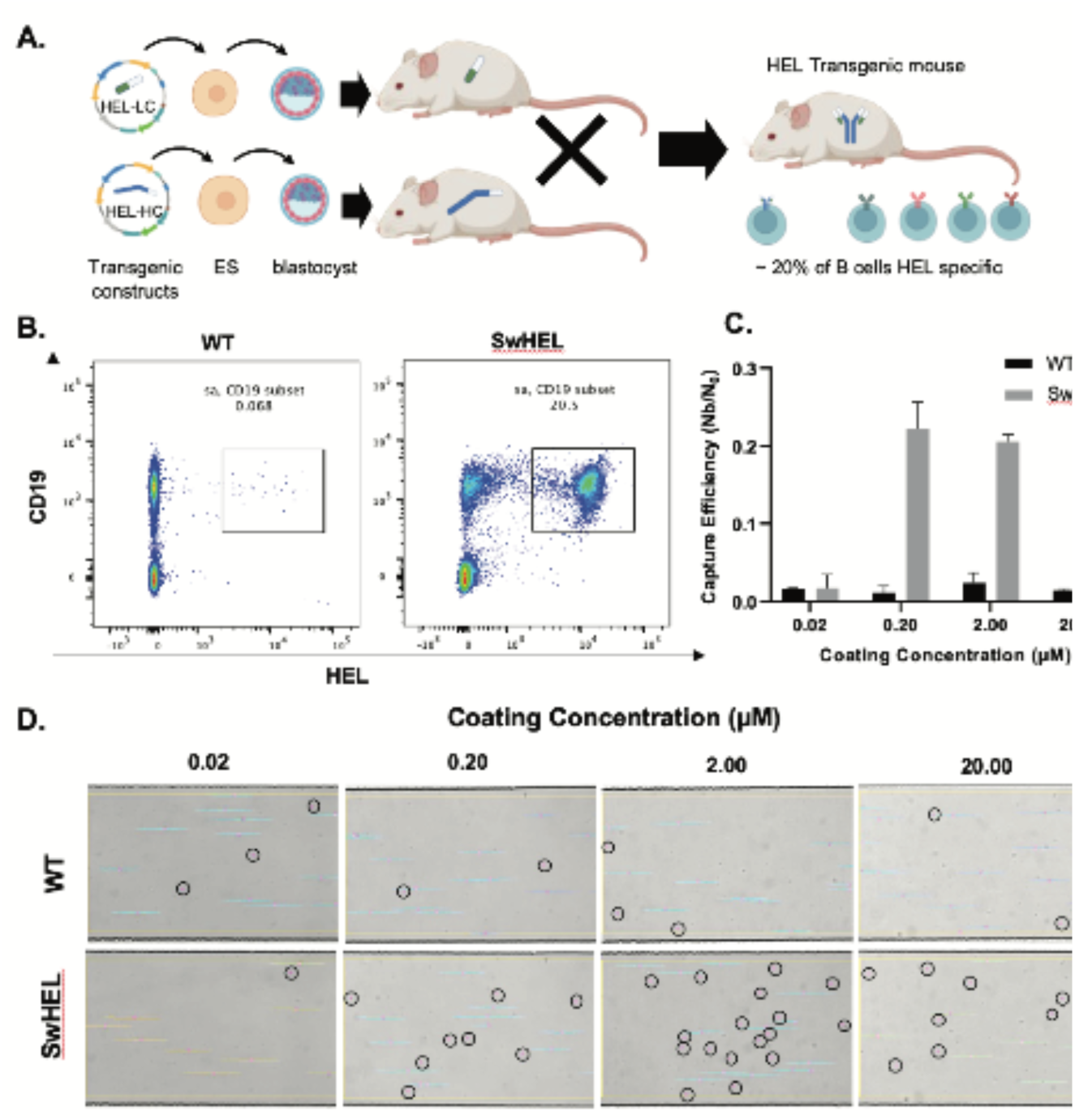
A) Schematic demonstrating how the SwHEL mouse is created such that ∼20% of the cells from the spleen at HEL-specific B cells (produced in part from BioRender). B) Flow cytometry showing subset of spleen cells that are specific to HEL from WT and SwHEL mice. C) Capture efficiency at 100 µL/h (n=3, mean ± SEM) of WT and SwHEL cells in microfluidic device coated with different concentrations of HEL and D) microscopic imaging showing cells under shear stress of 0.30 dyn/cm^2^ in the device. Circled cells are bound to HEL.

In peripheral blood of humans vaccinated for or infected by the influenza virus, circulating antibodies have a broad range of binding affinities for hemagglutinin (HA) ^30^, which is the major surface protein of the influenza A virus and is essential to the entry of the virus into host cells. To capture this range of antibody affinities, we used five single-cell hybridoma clones (Fig. 3A) that secrete monoclonal IgG antibodies specific to influenza A/Puerto Rico/8/34 virus HA (Table 1). Before measuring the binding kinetics of the membrane-bound BCR to HA in the microfluidic device, we first assessed the binding of the secreted monoclonal antibodies (mAbs) to both mammalian-expressed recombinant HA (affinity) and purified virus particles (avidity) using two well established methods: enzyme-linked immunoassay (ELISA) and oblique-incidence reflectivity difference (OI-RD) ^28, 31^. For ELISA, antibodies were serially diluted two-fold to generate antibody-binding curves to the virion. The binding curve was then fitted to extract K_A_ according to first-order binding kinetics (Fig. 3B). mAb binding was dose-dependent, with mAb H143 requiring 100-fold higher starting antibody concentration due to poor binding to the virion compared to the other four mAb tested. The equilibrium affinity constant (K_A_, M^-1^) of the mAbs to the virus particles varied more than 10-fold from 1 x 10^8^ M^-1^ to 1.5 x 10^9^ M^-1^ with the mAb H36 showing the highest K_A_, followed by mAb H163, H37, H35, and H143 (Fig. 3B).

**Figure 3.**
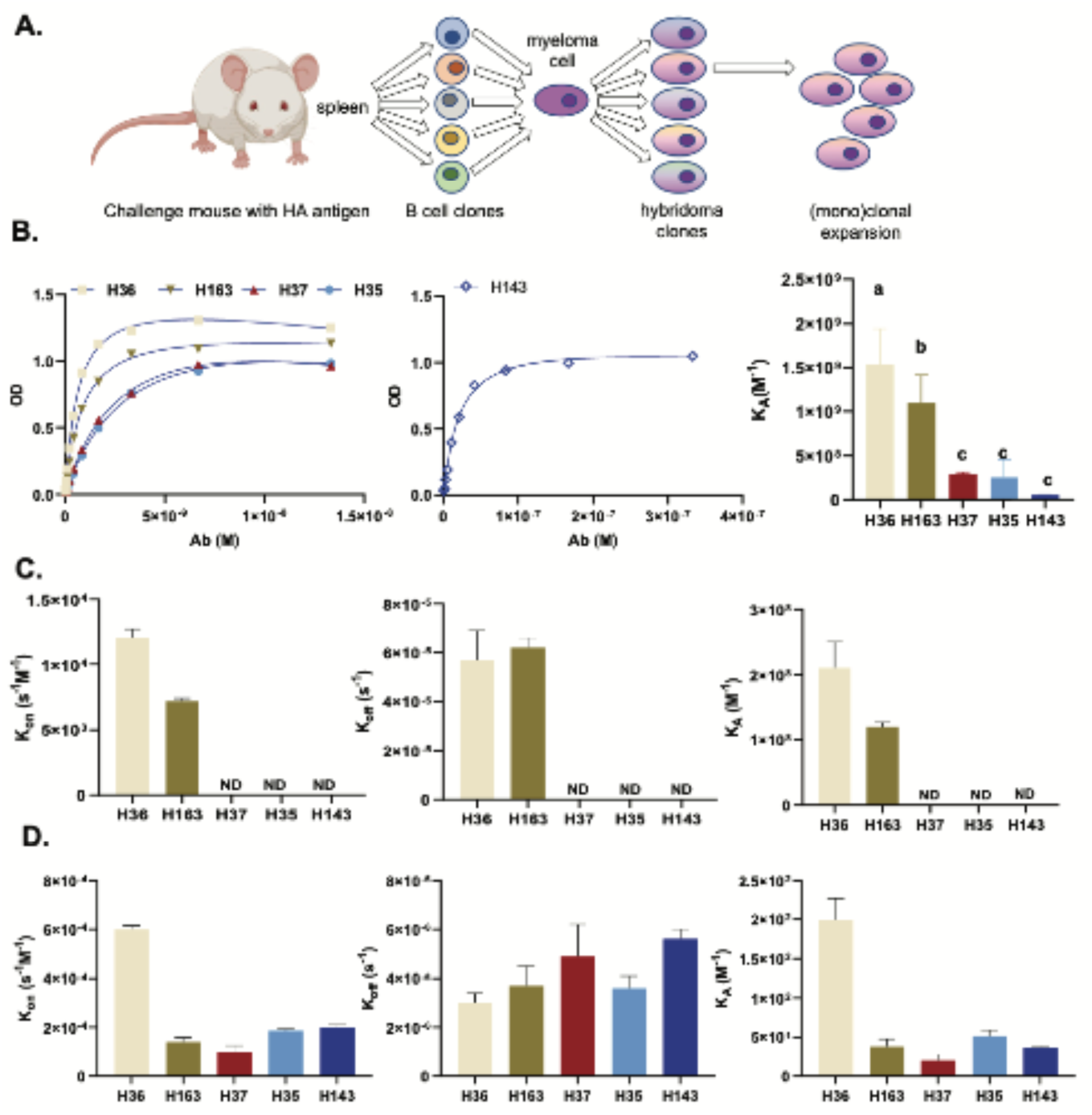
A) Schematic demonstrating how hybridoma technology produces monoclonal hybridoma clones (produced in part from BioRender). B) Optical Density binding curve generated by ELISA assay fitted to measure antibody affinity. C) Antibody affinity to HA and D) avidity to influenza virion measured by OI-RD.

The affinity of the antibodies to HA was also measured by OI-RD, where the antigen was circulated over fixed antibodies, then washed away to generate binding curves (Fig. S2). Similar to ELISA, H36 displayed the highest antibody affinity to HA, followed by H163 (Fig. 3C). The difference in equilibrium affinity between H36 and H163 was due to a higher 𝑘_*on*_ of H36 (Fig. 3C). The association and dissociation rates of antibodies from all other cell lines were below the detection threshold. Thus, the antibodies are considered to have low affinity to HA. Avidity measurements were similarly performed by OI-RD, but with the influenza virion immobilized instead of the HA-antigen. While H36 has the highest avidity (highest 𝑘_*on*_ and K_A_), all antibodies bound to the virion (Fig. 3D), further confirming the specificity of the antibodies to HA (Figs. 3A, D).

To observe how B cells (i.e. the membrane bound BCR) bind to HA in the device under force, cells from the five hybridoma lines and DS.1 (negative control) were perfused into a laminar microfluidic channel coated with HA (0.2 µM) under a range of constant shear force at the surface (0.03 – 0.15 dyne/cm^2^). Cells were perfused in the device at three flow rates (10, 20, and 50 μl/h), corresponding to three shear stresses at the surface (0.03, 0.06, and 0.15 dyne/cm^2^). These fluid flows are expected to apply 65, 130, and 325 pN forces which are in the physiological range of forces experienced by BCR-antigens bonds ^8^. The binding criterion was set to a minimum bound lifetime of 10, 20, 50, or 100 s.. For all binding criteria, H36 demonstrated the highest capture efficiency (Fig. 4A, B) ranging from as high as 0.78 (78%) at the lowest shear (0.03 dyne/cm^2^) and lowest minimum binding criteria (10 s), to as low as 0.08 (8%) at the highest shear (0.15 dyne/cm^2^) and highest minimum binding criteria (100 s). At the two higher shear rates, the other cell lines showed little binding in the device, regardless of binding criteria. At the lowest shear and all minimum binding criteria, H36 is followed by H163, H37, and H35 with H143 showing little to no binding.

**Figure 4.**
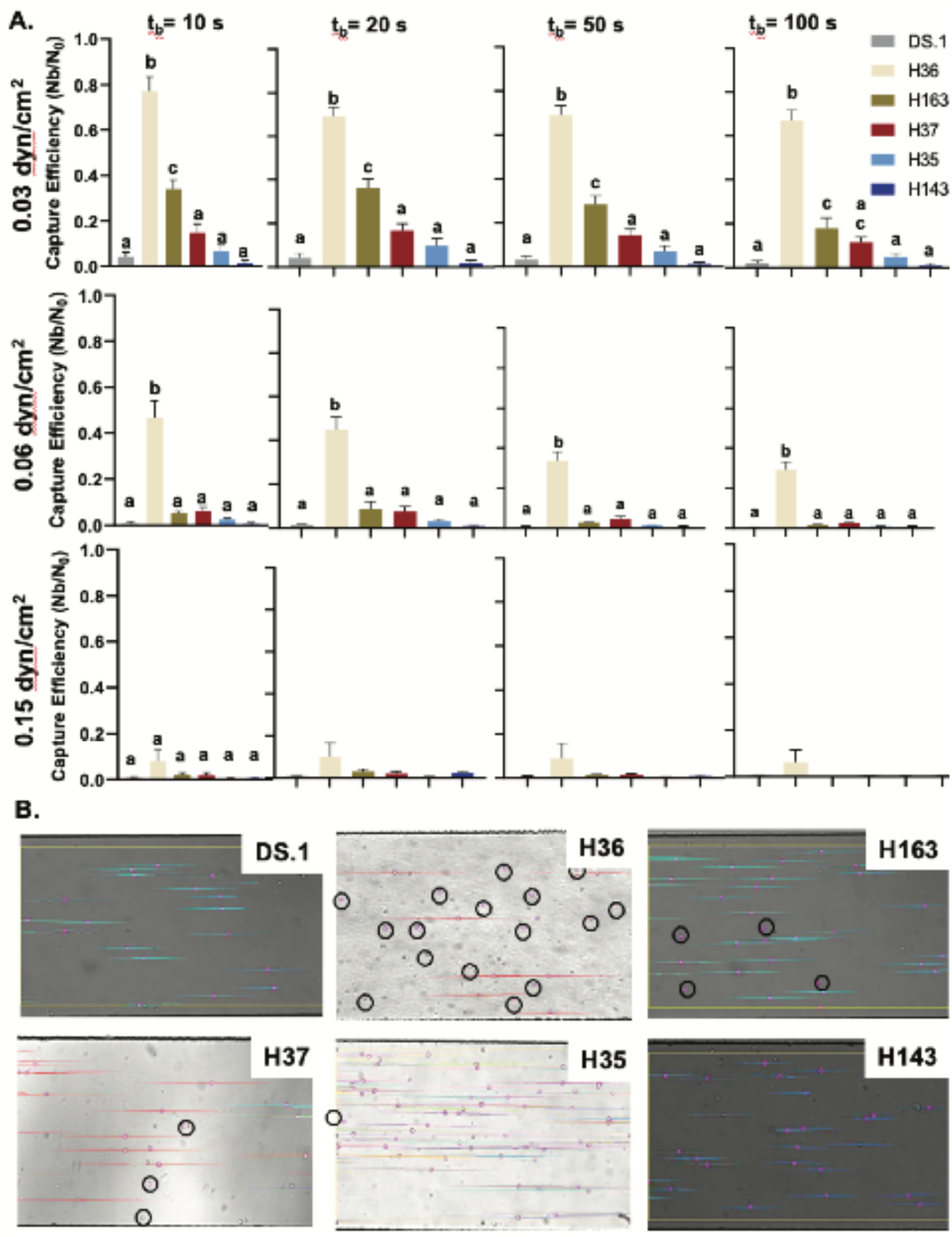
A) Capture efficiency (n ≥ 3, ± SEM) of five HA-specific hybridoma lines and negative control under shear stress of 0.03, 0.06 and 0.15 dyn/cm^2^ with minimum binding criteria of 10, 20, 50 and 100 s. B) Microscopic imaging showing cells under shear stress of 0.06 dyn/cm^2^ in the device. Circled cells are bound to HA.

The higher capture efficiency of H36 provides an opportunity to use the microfluidic device to enrich (or capture) a mixed population of hybridoma clones in H36. We created a mixed population of the five hybridoma cell lines as well as DS.1, with the concentration of H36 set at <5% representing a relatively dilute or rare cell (Fig. 5A). The three highest binding clones were fluorescently labeled with different colors, and the remaining three cell lines labeled with a fourth color. The mixed cell population was introduced into the device at two flows (10 and 20 μl/h) corresponding to the two lowest shear (0.03 and 0.06 dyne/cm^2^). Over 20% of the cells captured on the device (Fig. 5B-G) were H36, representing a 4-to-5-fold enrichment. H163 and H37 were neither enriched nor diluted on the surface, whereas the remaining three cell lines were diluted.

**Figure 5.**
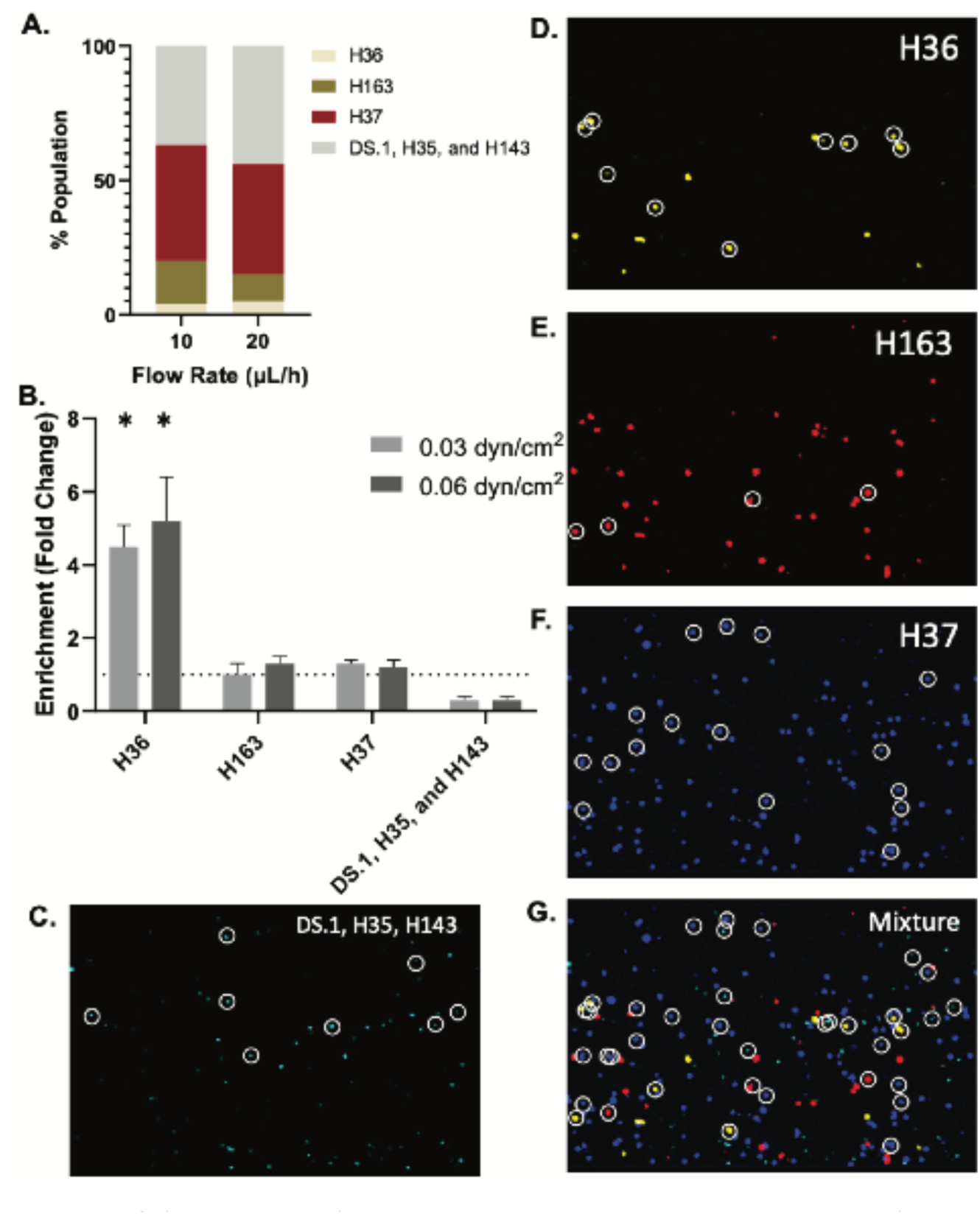
A) Composition of hybridoma cell lines in the mixture that is perfused in the device. B) Fold change in enrichment (Bound Cells (%)/Perfusate (%)) for different cell populations in mixture when perfused at 10 and 20 µL/h (n=3, mean ±SEM). C-G) Microscopic imaging showing cells under shear stress of 0.06 dyn/cm^2^. H36 (yellow), H163 (red), H37 (blue) and DS.1, H35 and H143 (cyan) are shown individually and in the mixture in which they were perfused. Circled cells are bound to HA. Bound cells were measured according to a minimum binding criterion of 10 seconds. Values that differ from 1-fold change with statistical significance (p < 0.05) are indicated.

BCR-antigen kinetic parameters are generally measured using the secreted antibodies. Here, we used membrane-bound BCR binding to HA under force to measure the reactive compliance (𝑥_β_) and effective 𝑘_*on*_and 𝑘_*off*_. To measure 𝑘_*on*_, a first order kinetic model was derived to fit cell binding data that depend on the number cells that experience an initial tethering of at least 1 s to the surface (Fig. 6A). Effective 𝑘_*on*_measurements for H143 and H35 were significantly lower than H36, H163 and H37 (Fig. 6B). To measure effective 𝑘_*off*_ and 𝑥_β_, force-dependent survival curves were generated for cells with minimum binding criteria of 10 s (Fig. 6C). Force-dependent values of 𝑘_*off*_ were then fitted to determine 𝑥_β_. The fit was extrapolated to zero-force to determine effective 𝑘_*off*_ (Fig. 6D). Effective 𝑘_*off*_ is lowest for H36 and H37. While H163 has a 𝑘_*on*_ similar to that of H36, its 𝑘_*off*_ is significantly higher (Fig. 6E). Additionally, 𝑥_β_measurements show that not all hybridoma cells behave the same when subjected to shear force in the device, with H35 and H143 having lower 𝑥_β_ compared to H37, consistent with off-rates that are less sensitive to force (Fig. 6F). Finally, the affinity constant, the ratio of the effective on-and off-rates, shows that H36 has the highest affinity to HA (Fig. 6G). Comparing these measurements in the device to the affinity measurements by ELISA and OI-RD shows a strong correlation between all three assays (Fig. S3).

**Figure 6.**
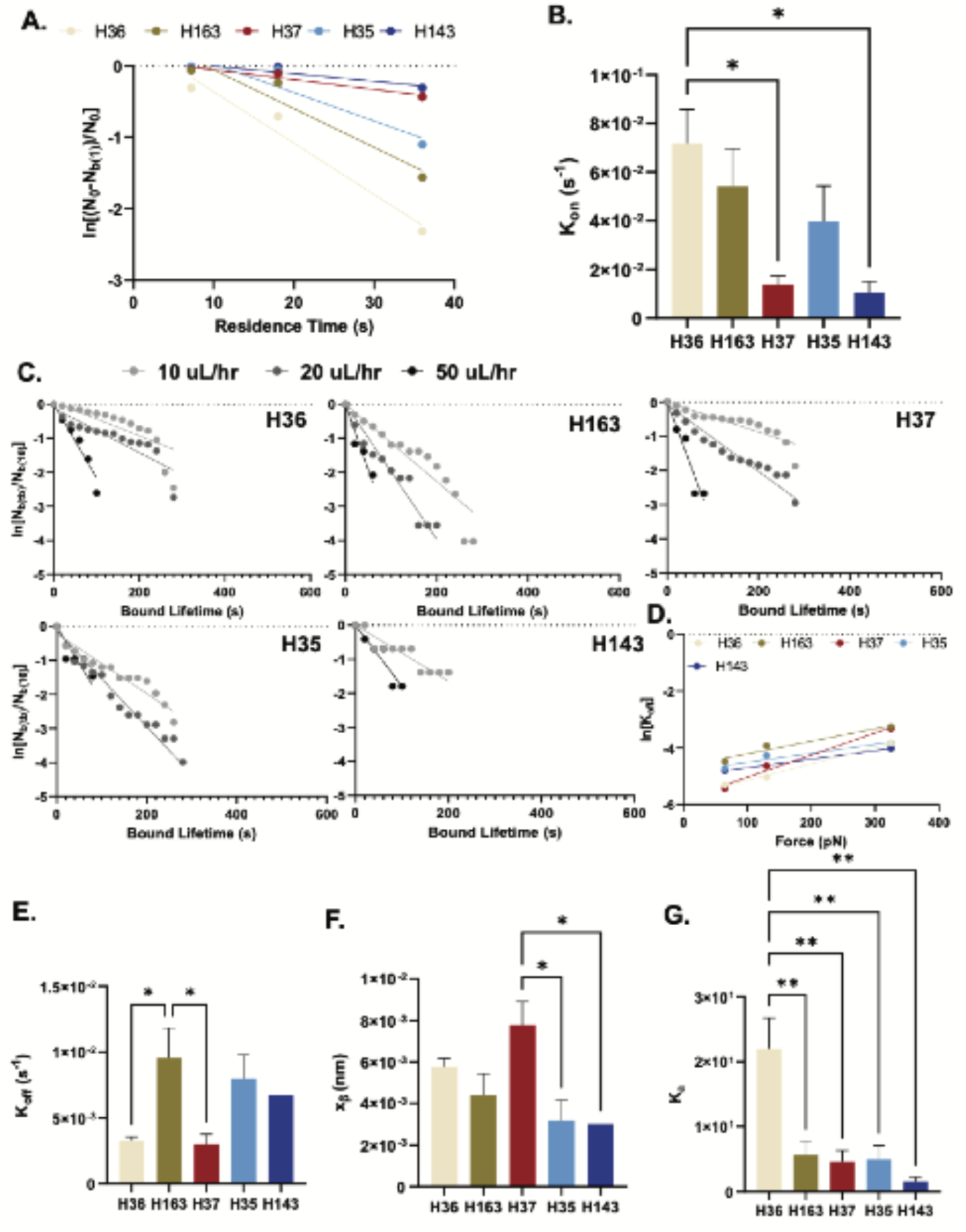
A) Data fit based on first-order kinetic model to measure effective 𝑘_*on*_. B) Measured values of effective 𝑘_*on*_ for the different hybridoma cell lines. C) Survival curve fits to determine force-dependent 𝑘_*off*_, where bound lifetime is the length of time the cells remain bound for more than 10s and D) data fit to determine reactive compliance (𝑥_β_) and effective 𝑘_*off*_ at zero-force. E) Values of effective 𝑘_*off*_ at zero-force and F) reactive compliance (𝑥_β_). G) Membrane-bound BCR affinity (effective 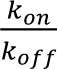) to HA

## DISCUSSION

Recent efforts to optimize isolation of high affinity antibodies for therapeutic applications or to assess the functional immune status of individuals have highlighted the importance of rapid screening and characterization of B cells and their antibodies ^32–38^. These methods rely on equilibrium binding properties of BCR/antibody-antigen bonds, followed by cumbersome and expensive steps of cloning, expression, and mAb purification. Using a very simple microfluidic device (single rectangular channel), we demonstrate that by flowing B cells at a controlled shear over a surface functionalized with antigen, we can rapidly and efficiently capture and enrich a population of antigen-specific high affinity B cells. Furthermore, this methodology can also be used to assess kinetic binding parameters of the membrane-bound BCR (𝑘_*on*_, 𝑘_*off*_, 𝐾_5,_ and 𝑥_β_). We found that 𝐾_5_ correlated well with traditional assays that measure the equilibrium binding affinity of the secreted antibody. Our results demonstrate that a simple microfluidic strategy can rapidly identify antigen-specific, high affinity, rare B cells from a larger population, and thus could be used as a cost-effective strategy to identify B cell clones generating mAb for a host of clinical applications or rapidly assess the functional immune status of an individual.

The creation of antigen-specific high affinity mAbs has a host of important clinical applications including the treatment of patients with viral infections (most recently COVID-19 ^39^), allergic inflammation in diseases such as asthma ^40^, autoimmune diseases such as rheumatoid arthritis ^41^, and cancer ^42^. All told, the annual global sales of therapeutic and prophylactic mAbs is in excess of US$75 billion ^43^. The process to create mAbs utilizes either hybridoma technology or antibody phage display ^44^. Although generally effective, both techniques have significant drawbacks, including the cost and overall time (months to years once an antigen has been identified) to create the mAb ^45^. This is particularly important for creation of mAbs for diseases with a rapidly changing antigen landscape such as during infections with highly mutating RNA viruses (e.g., omicron variant of SARS-CoV2 ^46^) and most solid cancers ^47^. A key result of our study is the high correlation between the affinity of the membrane-bound BCR measured in our microfluidic device and the affinity of the secreted antibodies measured by ELISA and OI-RD. As such, our microfluidic strategy, which employs tunable force-dependent (shear stress) binding, can be used to capture and enrich very high affinity B cell clones from a mixed population, and thus the sources of antigen-specific high affinity mAbs. Using only a single pass at a single flow (shear stress), we enriched a dilute (<5%) population of high affinity B cells by 4-to-5 fold. One can then easily imagine repeating this process to further enrich, or using higher flows to capture extremely high affinity clones from a larger population of B cells. The distinct advantage of this approach over current methods is it obviates the laborious steps of antibody testing and screening because the force-dependent selection of the high affinity B cell clone has already performed this task.

An alternative to our microfluidic strategy to identify antigen-specific, high affinity B cell clones is to utilize affinity maturation ^48^. Following exposure to a pathogen, B cells undergo affinity maturation in which somatic hypermutation creates BCRs with higher antigen affinity and thus improved immune response to a specific antigen. Affinity maturation creates a diverse population of B cells with frequently changing characteristics ^49^. Although BCR affinity is thought to predict functional immunity ^4^, what features of the diverse population of B cells are predictive is not known. For example, does the high affinity B cells predict functional immunity? If so, how high is “high”? Alternatively, a population of lower affinity B cells may be adequate. If this is the case, what range of affinity is required? By demonstrating that our technology can capture and enrich a high affinity B cell subpopulation, it is easy to extrapolate how the technology could be used to quantify the full spectrum of B cell binding affinity to a target antigen. For example, the first step (high shear and potentially multiple passes) captures the highest affinity B cells in the device, which can then easily be removed and collected. The population of B cells which remain (unbound from the first pass) are collected and passed through the device at a lower shear. The next highest affinity B cells attach to the surface, which can then be collected. The cells which did not bind, are collected and the process repeated at a lower shear. In this fashion, a series of B cell populations are collected and quantified in a progressively descending order of affinity. Although most of our studies were generated using B cell lines, in proof-of-concept studies with primary spleen cells from SwHEL BCR transgenic mice, expressing a BCR of known high affinity for HEL ^15^, we demonstrate that our device also allows study of primary cells. Assessing a population of B cells quantitatively and at an individual level reliably and affordably could be instrumental in pandemic responses. For example, rapidly assessing the spectrum of B cell binding affinity at a point in time could provide a surrogate for the immune status of individuals, and thus provide important information for the strategic distribution of limited resources (drug and vaccine).

In summary, we present a simple microfluidic strategy that utilizes shear stress (force) to characterize the force-dependent antigen-specific binding characteristics (𝑘_*on*_, 𝑘_*off*_, 𝐾_*A*_, and 𝑥_β_) of the membrane-bound BCR. We show that the binding affinity of five hybridoma cell lines specific to influenza HA is remarkably variable, but that the affinity of the cell membrane-bound BCR correlates well with the binding affinity of the secreted antibodies. The technology can be used to easily capture and enrich a population of antigen-specific, high affinity B cells, and thus be used to quantitatively characterize the full spectrum of binding affinity for a diverse population of B cells. The technology is easily scalable and thus has potentially important applications to simplify and reduce the cost and time to create mAb, as well as to rapidly and cost-effectively assess the spectrum of B cell affinity and thus the functional immune status of an individual.

## ACKNOWLEDGEMENTS

We would like to thank UC Davis Center for Nano and Micro Manufacturing (CNM2) for assistance in the microfabrication. This work was supported by the National Institutes of Health (R21 AI161041).

## SUPPLEMENTAL INFORMATION

**Figure S1.**
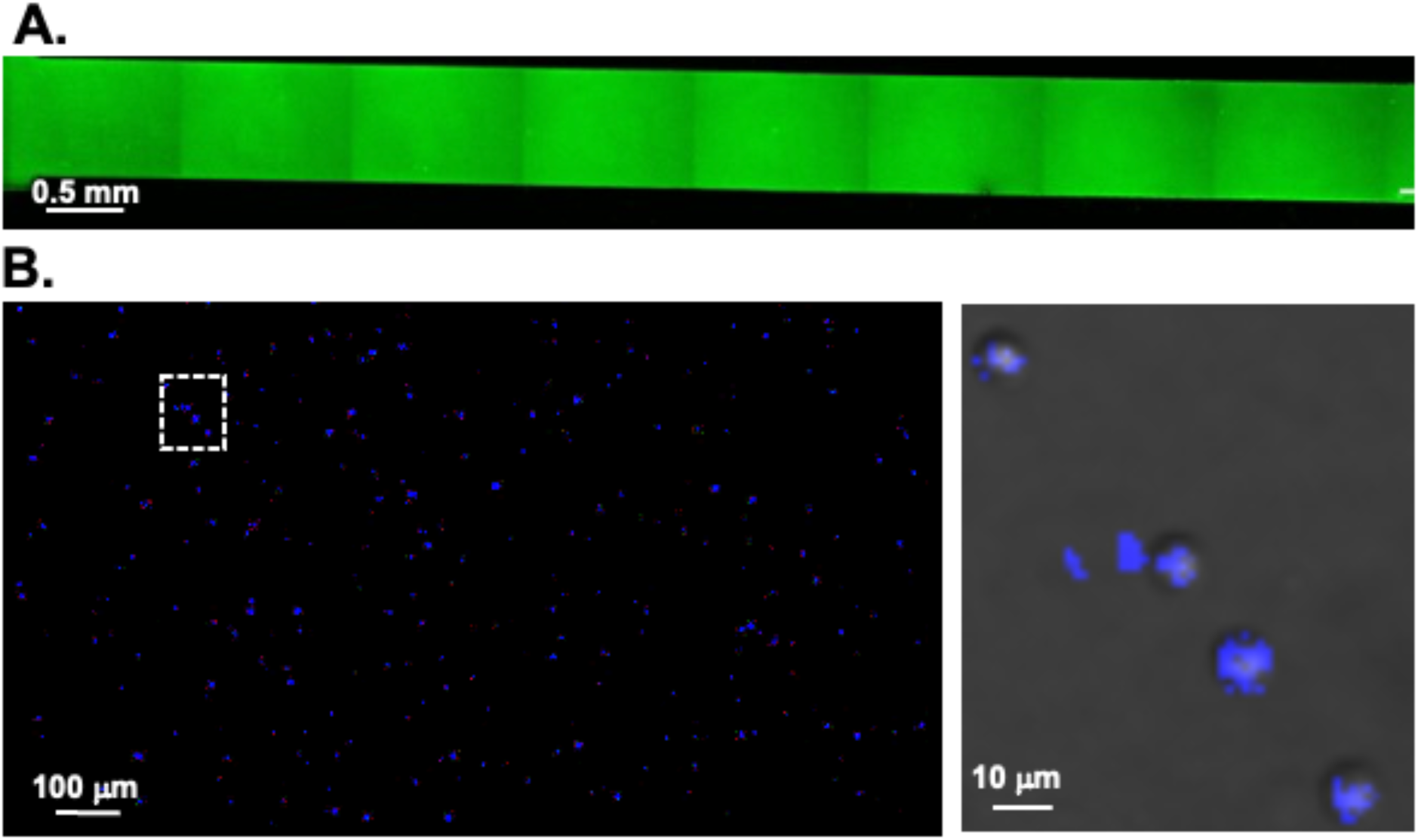
Biotin-FITC bound to NeutrAvidin coated device (A). CD19 coated SwHEL cells bound to HEL in device (B).

**Figure S2.**
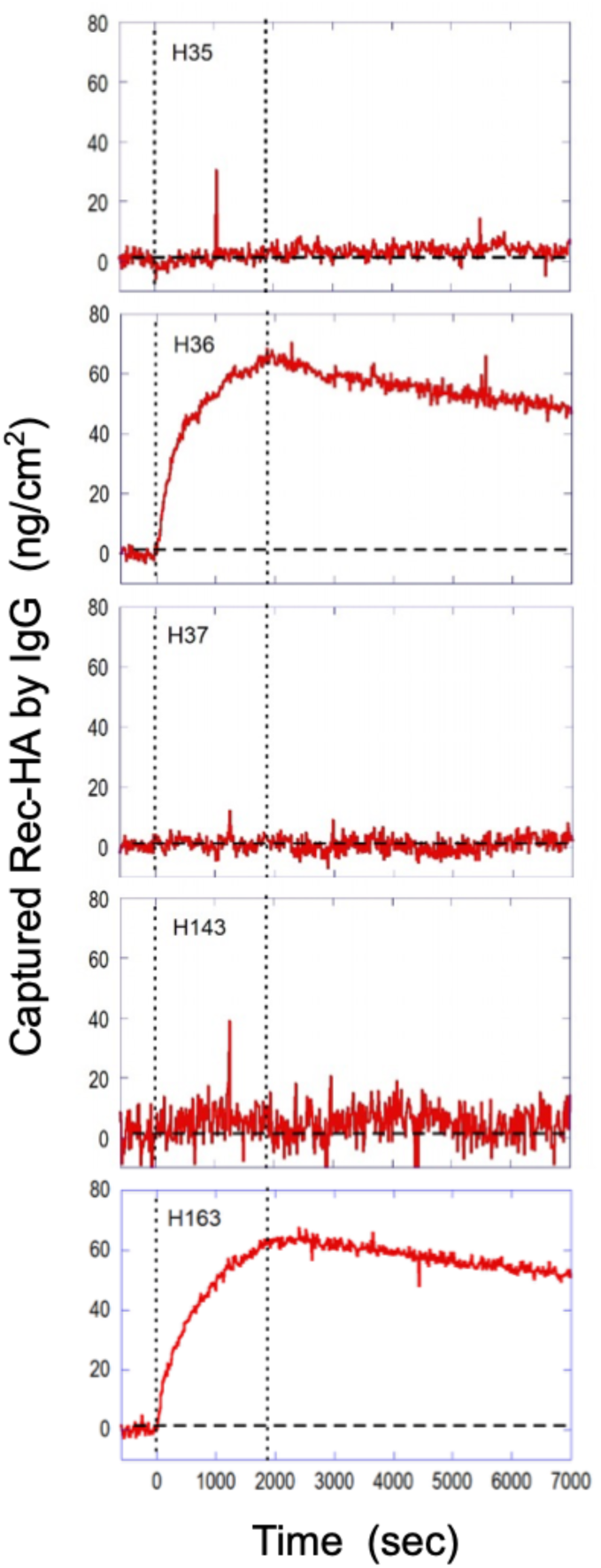
Binding curves of Recombinant HA (Rec-HA) to H35, H36, H37, H143, and H163 using an OI-RD detected, microarray-based assay system. Two vertical dotted lines mark starts of association phase and dissociation phase of the affinity assay. The concentration of the recombinant HA is 300 nM. The curves are fit to a 1-to-1 Langmuir reaction model to yield association rate constants k_on_ (M^-1^sec^-1^) and dissociation rate constants k_off_ (sec^-1^). Affinity constants K_a_ of Rec-HA to IgG are determined from K_a_ = k_on_ / k_off_.

**Figure S3.**
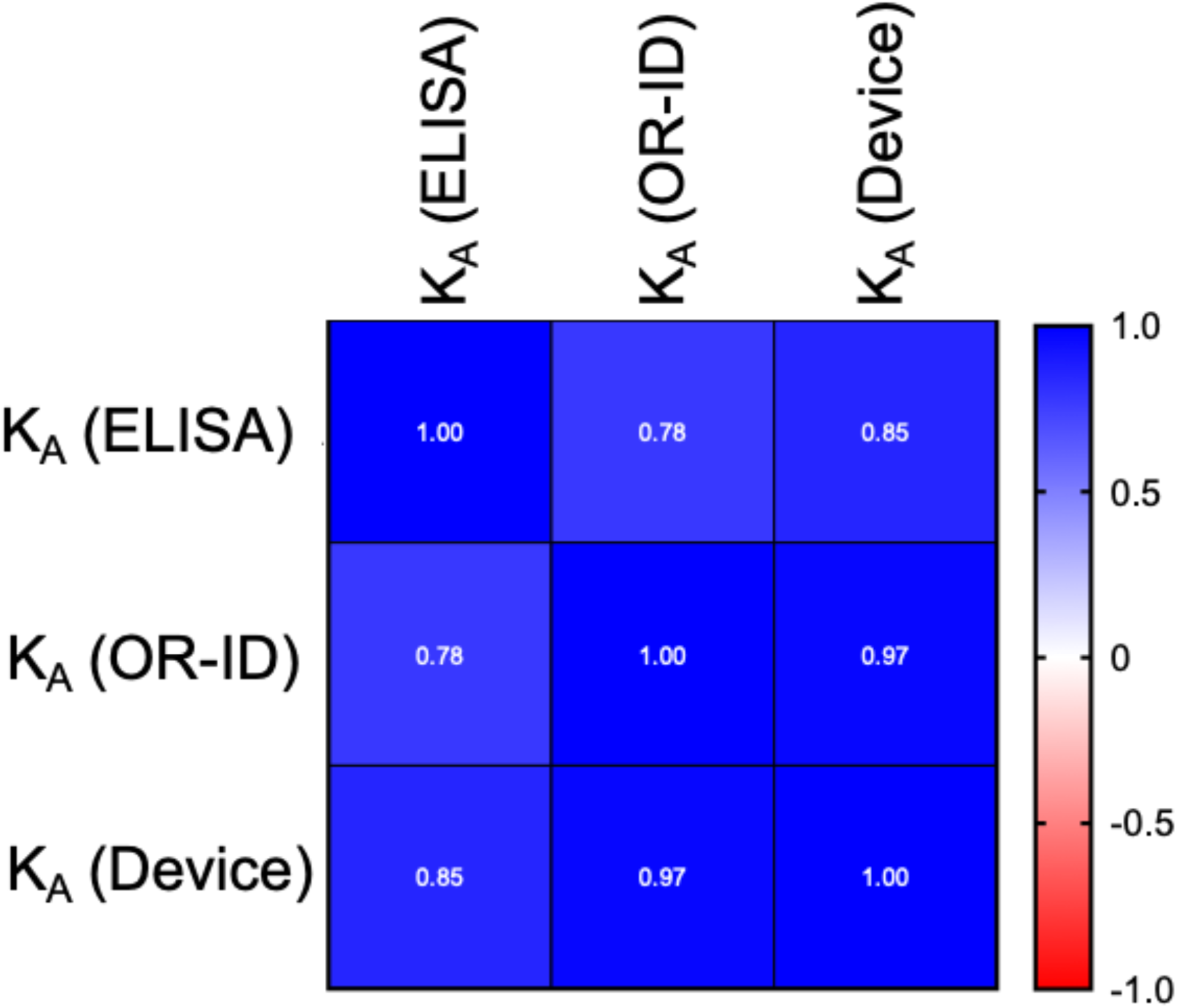
Pearson correlation matrix of K_A_ values measured by three different assays: ELISA, OI-RD and microfluidic device.

